# DNA Methylation-based Signatures Classify Sporadic Pituitary Tumors According to Clinicopathological Features

**DOI:** 10.1101/2020.04.25.061903

**Authors:** Maritza S. Mosella, Thais S. Sabedot, Tiago C. Silva, Tathiane M. Malta, Felipe D. Segato, Karam P. Asmaro, Michael Wells, Abir Mukherjee, Laila M. Poisson, James Snyder, Ana C. deCarvalho, Tobias Walbert T, Todd Aho, Steven Kalkanis, Paula C. Elias, Sonir R. Antonini, Jack Rock, Houtan Noushmehr, Margaret Castro, Ana Valeria Castro

**Affiliations:** Department of Neurosurgery, Hermelin Brain Tumor Center, Henry Ford Health System, Detroit, MI, USA; Department of Neurosurgery, Omics Laboratory, Henry Ford Health System, Detroit, MI, USA; Department of Genetics, Ribeirao Preto Medical School, University of São Paulo, Ribeirao Preto, Brazil; Department of Pathology, Henry Ford Health System, Detroit, MI, USA; Department of Biostatistics, Henry Ford Health System, Detroit, MI, USA; Department of Radiology, Henry Ford Health System, Detroit, MI, USA; Internal Medicine Department, Ribeirao Preto Medical School, University of São Paulo, Ribeirao Preto, Brazil; Department of Pediatrics, Ribeirao Preto Medical School, University of São Paulo, Ribeirao Preto, Brazil

**Author notes:** Corresponding author: Ana Valeria Castro, Scientist/Professor, Department of Neurosurgery, Hermelin Brain Tumor Center, Henry Ford Health System, 2799 West Grand Blvd, E&R 3096, Detroit, MI, 48202. University of Miami, FL, USA. University of São Paulo, Ribeirao Preto, Brazil (Tathiane M. Malta). Contributed equally (Houtan Noushmehr, Margaret Castro, Ana Valeria Castro). **Authorship:** Overall concept and coordination of the study: AVC, MC, HN, SA, PE; retrieval of publicly available molecular and clinical data: MSM; Bioinformatic and statistical analyses: MSM, TSS, TCS, TMM, FSD, HN and input from LMP; HFHF cohort: pathology review MF, AM; clinicopathological and radiological data were generation and collection: AVC, MSM, KPA, AM, MF, TA; molecular data generation: AC, TMM; manuscript was written by MSM, AVC, HN, intellectual contribution: JS, SK, TW. All authors contributed to the revision of the manuscript.

**Keywords:** DNA methylation, classification, regulatory elements, enhancers, pituitary tumors

## Abstract

**Background:** Distinct genome-wide methylation patterns have consistently clustered pituitary neuroendocrine tumors (PT) into molecular groups associated with specific clinicopathological features. Here we aim to identify, characterize and validate the methylation signatures that objectively classify PT into those molecular groups.

**Methods:** Combining *in-house* and publicly available data, we conducted an analysis of the methylome profile of a comprehensive cohort of 177 tumor and 20 non-tumor specimens from the pituitary gland. We also retrieved methylome data from an independent pituitary tumor (PT) cohort (N=86) to validate our findings.

**Results:** We identified three methylation clusters associated with functional status and adenohypophyseal cell lineages using an unsupervised approach. We also identified signatures based on differentially methylated CpG probes (DMP), some of which overlapped with pituitary-specific transcription factors genes (SF1 and Tpit), that significantly distinguished pairs of clusters related to functional status and adenohypophyseal cell lineage. These findings were reproduced in an independent cohort, validating these methylation signatures. The DMPs were mainly annotated in enhancer regions associated with pathways and genes involved in cell identity and tumorigenesis.

**Conclusions:** We identified and validated methylation signatures that distinguished PT by distinct functional status and adenohypophyseal cell lineages. These signatures, annotated in enhancer regions, indicate the importance of these elements in pituitary tumorigenesis. They also provide an unbiased approach to classify pituitary tumors according to the most recent classification recommended by the WHO 2017 using methylation profiling.

**Key-points:**

- Distinct methylation landscapes define PT groups with specific functional status/subtypes and adenohypophyseal lineages subtypes.
- Methylation abnormalities in each cluster mainly occur in CpG annotated in distal regions overlapping predicted enhancers regions associated with pathways and genes involved in cell identity and tumorigenesis.
- DNA methylation signatures provide an unbiased approach to classify PT.

**Importance of the study:** This study harnessed the largest methylome data to date from a comprehensive cohort of pituitary specimens obtained from four different institutions. We identified and validated methylation signatures that distinguished pituitary tumors into molecular groups that reflect the functionality and adenohypophyseal cell lineages of these tumors. These signatures, mainly located in enhancers, are associated with pathways and genes involved in cell identity and tumorigenesis. Our results show that methylome profiling provides an objective approach to classify PT according to the most recent classification of PT recommended by the 2017 WHO.

## 1 Introduction

Pituitary tumors (PT), a type of neuroendocrine tumors, comprise the second most common neoplasm of the central nervous system (CNS) (∼17%) ^1–3^ with an annual average age-adjusted incidence of 4.08 per 100,000 population ^3^. Stratified by endocrine status, PT are classified as functioning (FPT) (46-64%) or nonfunctioning (NFPT) (36%–54%) subtypes, the latter also referred to as silent adenomas ^1,4^. In relation to the histopathological classification the most recent recommendation by the World Health Organization (WHO) in 2017 is based on immunostaining for adenohypophyseal hormones and cell-lineage ^5^. Despite its importance and extensive use for tumor classification, the immunopathology method is limited by its subjectivity and inaccuracy in pituitary and several other tumors ^6,7^. To circumvent these limitations and/or to complement pathology results, inclusion of genomic-based signatures such as recurrent somatic mutations and chromosome number alterations has advanced and refined the classification of many tumors, including CNS neoplasms ^8,9^. However, in contrast to other CNS tumors, genomic alterations are rare in PT or are restricted to some subtypes (*i.e*. GNAS mutation in somatotroph adenomas) ^10–12^. Contrariwise, distinct genome-wide DNA methylation patterns are consistently associated with their clinicopathological features such as function and histological variants. In contrast to other CNS tumors, the importance of methylation markers in the taxonomy of PT has not been explored to date ^7,10,13–17^. A recent study reported that the combination of genome-wide molecular profiling of multiple platforms, including DNA methylation, provided an objective approach to the classification of PT ^10^. However, the identification of specific methylation signatures to classify PT into these methylome-based groups has not been exploited.

To address this knowledge gap, we aimed to identify, characterize and validate methylation-based markers that define PT according to clinicopathological features. For this purpose, we compiled and performed a comprehensive analysis of the genome-wide DNA methylation data on 177 adult sporadic tumors and 20 non-tumorous tissue from the pituitary gland obtained from a multicenter source and validated our findings in an independent cohort. We identified methylation signatures that classified PT according to methylation groups that reflected the functionality and the adenohypophyseal lineages of these tumors.

## 2 Subjects and methods

### 2.1 Patients and tissue specimens

DNA methylation data were retrieved from 3 publicly available cohort datasets (data retrieval freeze on August 2018) ^7,13,14^ and from our cohort at the Hermelin Brain Tumor Center (HBTC) of the Henry Ford Health System (Detroit, MI, USA) (n=23; HBTC cohort, unpublished). The final cohort consists of 20 non-neoplastic pituitary glands (non-tumor/NT) and 177 PT from patients over 18 years (Pan-pit), composed of 87 FPT and 90 NFPT. Clinical information about this study cohort is summarized in Table 1 and detailed in Supplementary Table S1. The PT clinicopathological classification criteria of features such as invasion from our internal cohort were matched with the criteria provided by the authors of the external cohorts ^7,13,14^. Specific transcription factor-based adenohypophyseal lineages were assigned to each PT subtype in accordance with the 2017 WHO classification ^5^. All the studies were approved by their respective Institutional Review Boards and ethics committees and written consent obtained from each patient at the attending institutions ^7,10,13,14^.

### 2.2 Data retrieval and preprocessing

DNA from pituitary tissue was obtained from fresh-frozen (FF) ^13,14^ or formalin-fixed-paraffin-embedded specimens (FFPE) ^7^. The method of DNA extraction, processing, and assays for each cohort included in this study is reported in the respective manuscripts ^7,13,14^ and for the HBTC cohort is available in the Supplementary Methods. The IDAT files for the DNA methylation bead arrays (450K or 850K/EPIC) from the Ling, Kober, and Capper cohorts ^7,13,14^, were retrieved through the GEOquery package ^18^ and preprocessed simultaneously using minfi package^19^. The EPIC array data from Capper and HBTC were processed separately using the same approach. Detailed information about preprocessing of IDAT files is depicted in Supplementary methods. Principal component analysis (PCA) revealed that there were no batch effects related to the cohort, tissue sources (FFPE or FF) and array platform (450K/EPIC) (Supplementary Figure S1A-C). The HBTC DNA methylation data was submitted to Gene Expression Omnibus (GSEXXXX).

To test the reproducibility of our findings, we retrieved the methylome data from 86 patients with PT obtained from an independent study published after our data freeze (EMBL-EBI accession number E-MTAB-7762) ^10^.

### 2.3 Unsupervised and supervised analysis of the PT methylome

To reduce noise, the most variable probes were sorted according to standard deviation across the 177 PT specimens. Next, a consensus clustering analysis^20^ was applied to the top 1% most variant probes (n=4,526). Agglomerative hierarchical clustering with 1000 resampling steps, upon euclidean distance was used, as described previously ^8,20^. Optimal clusters were selected based on statistical parameters: Calinski-Harabasz curve and consensus cluster delta area and overlapped with clinicopathological annotations.

Two-tailed Wilcoxon rank sum tests were conducted for each CpG in order to identify pairwise differentially methylated probes (DMP) between the methylation clusters and invasion status. DMP were defined based on differences in mean methylation (diffmean) and adjusted p-value significance. For those comparisons with the highest number of significant probes (*e.g.* for NFPT-e *vs*. FPT-e), we applied a more stringent adjusted p-value cutoff, in order to retrieve DMP sets with similar number of probes. An absolute diffmean of 0.3 was used as a threshold for all methylation clusters comparisons. Between ACTH-e *vs.* NFPT-e clusters comparison adjusted p-values <1e-17 were obtained for both hypo- and hypermethylated probes; for ACTH-e *vs.* FPT-e and NFPT-e *vs.* FPT-e comparisons, the following adjusted p-values for the respective hypo and hypermethylated probes were obtained: p< 1e-16 and p<1e-19 and p<1e-21 p<1e-38, respectively. No significant adjusted p-values were obtained from the invasion status comparative analysis, therefore, the results are related to non-adjusted p-value <0.001 along with delta diffmean of 0.15, similar to the approach reported by other authors ^13,14^.

CpG probes were mapped to their CpG genomic location as CpG islands (CGI), shores, shelves, and open sea regions as previously defined ^21–24^. DMPs were integrated with predicted enhancers listed in the GeneHancer database ^25^ through the Genomic Ranges package (https://bioconductor.org/packages/release/bioc/html/GenomicRanges.html). Enrichment or depletion frequencies of DMP sets by genomic location and enhancer overlap were calculated as Odds Ratio using the 485,000 probes from HM450K platform as the reference.

Determination of the number of overlapping and distinct DMPs (hyper or hypomethylated) between pairwise comparisons were calculated using the UpsetR package ^26^.

### 2.4 DNA motif analysis

To predict transcription factor binding sites, we surveyed the DNA motifs within enhancer regions which overlapped our DMPs (namely enhancer-related DMP or eDMP). Sequence motifs were identified within ±200 bp from the core of the hyper- or hypomethylated enhancer regions using the bioinformatic tools HOMER ^27^. To increase the sensitivity of motif detection, up to two mismatches were allowed in each oligonucleotide sequence. The distributions of motif content in ‘target’ (*i.e.*, the enhancer regions) and random ‘background’ sequences were assessed for fold change values.

### 2.5 Integrative analysis between methylation and enhancer prediction databank

We identified CpG probes that overlapped with their predicted regulatory regions listed in the GeneHancer database ^25^. Based on the GeneHancer’s annotation confidence score for functional enhancers ^25^, the highest scored enhancers for each DMP set were selected. The selected enhancers were connected to their putative target genes as defined by four experimental metrics ^25^. The genes with the highest enhancer-gene pairing association score were selected (GeneHancer interaction score)^25^ and their expressions per methylation cluster are depicted as boxplots.

### 2.6 Methylation status of pituitary-specific transcription factors-related probes

We assessed the DNA methylation profile of CpG probes related to pituitary-specific transcription factor (TF) coding genes, known to determine the development of various adenohypophyseal cell types in the pituitary gland: Pit-1 (POU1F1), Tpit (TBX19), ER-alpha (ESR1), GATA2 and SF1 (NR5A1) ^1,22^. Next, we overlapped the pituitary-specific TF-related probes with the three DMP sets obtained from the pairwise comparison between the clusters. For each significant TF-related probe, we established methylation degree cutoffs that distinguished between clusters. The accuracy of these cutoffs in defining PT subtypes was investigated in the validation cohort ^10^.

## 3 Results

### 3.1 Genome-wide DNA methylation patterns define three groups of pituitary tumors related to specific functional subtypes and adenohypophyseal cell-lineages

Consensus clustering of the most variable DNA methylation probes in the pan-pit dataset, revealed three main clusters. The clusters were named according to their enrichment for specific functional subtypes (Figure 1A-B and Supplementary Figure S1C). The adrenocorticotropic hormone-enriched cluster (ACTH-e; n=29) is composed mainly of corticotroph adenomas (n=21) and 8 NFPT, including 1 silent corticotroph adenoma, 1 gonadotroph, 1 null cell adenoma and 5 of unknown histological NFPT subtypes. The NFPT-e cluster (n=82) consists mainly of NFPT (n=69), including 60 gonadotrophs, 7 null cell tumors, and 2 mixed non-functioning adenomas (LH/FSH/TSH), 1 FPT (lactotroph adenoma) and 12 tumors with unknown histological subtypes. The FPT-e cluster (n=66) is composed mainly of FPT (n=65), *i.e.*, 41 somatotrophs, 11 thyrotrophs, 8 lactotrophs, 5 plurihormonal/mixed adenomas (2 GH/PRL, 1 GH/TSH, 2 GH/PRL/TSH) and 1 NFPT (gonadotroph). The top most variant probes (according to standard deviation) across tumor specimens (n=4,526) are hypomethylated in the FPT-e cluster, hypermethylated in the NFPT-e cluster, and present an intermediate methylation pattern in the ACTH-e cluster (Figure 1B and 1C). ACTH-e, NFPT-e and FPT-e clusters are composed mainly by Tpit- (22/24; 92%); SF1- (60/70; 86%) and Pit-1 (65/66; 98%) cell lineages-derived subtypes, respectively.

**Figure 1.**
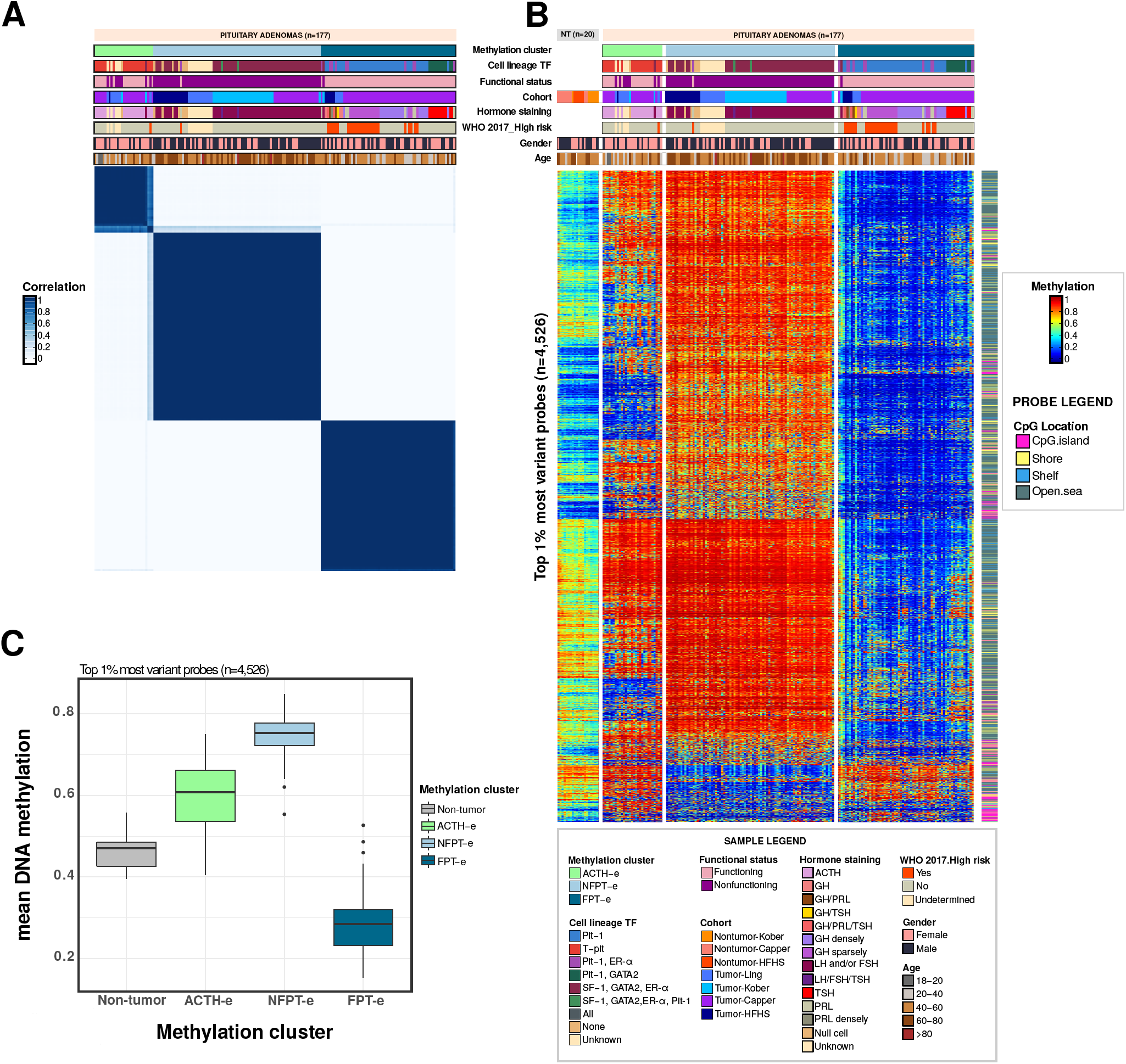
Methylome landscape segregates pituitary tumors into three distinct groups in a multicenter cohort. **(A)** Consensus Cluster correlation heatmap depicting three identified methylation-based groups among pituitary tumors. **(B)** Heatmap of the most variant CpG methylated probes (n=4,526) among 177 pituitary tumors according to their methylation-based cluster and for 20 non-tumor pituitary specimens. Columns represent specimens; rows represent the CpG probes. **(C)** Boxplots depicting the mean DNA methylation of the top most variant probes according to non-tumor and tumor pituitary specimens methylation groups.

### 3.2 Differential methylation between the three methylation-based PT clusters is observed in distal regions overlapping enhancer elements

The supervised comparisons between the clusters retrieved three DMP sets totaling 3,316 differentially methylated probes (Figure 2A discovery heatmap) that distinguished among the groups which were validated in an external cohort ^10^ (Figure 2A, validation heatmap). We observed that 1,973 (59%) out of 3,316 DMPs were located in enhancer regions (Figure 2A, left row label track). DMPs were enriched in open sea regions (n=2,257, 68%), while depleted in CGI, for the three pairwise methylation cluster comparisons (Figure 2B). Detailed information about the distribution of the number of DMPs according to methylation direction and CpG annotation is depicted in Table S2 (Supplemental material). The survey of common or exclusive DMPs among the methylation cluster comparisons are depicted in Figure 2C and D, based on the methylation direction (hypo or hypermethylated).

**Figure 2.**
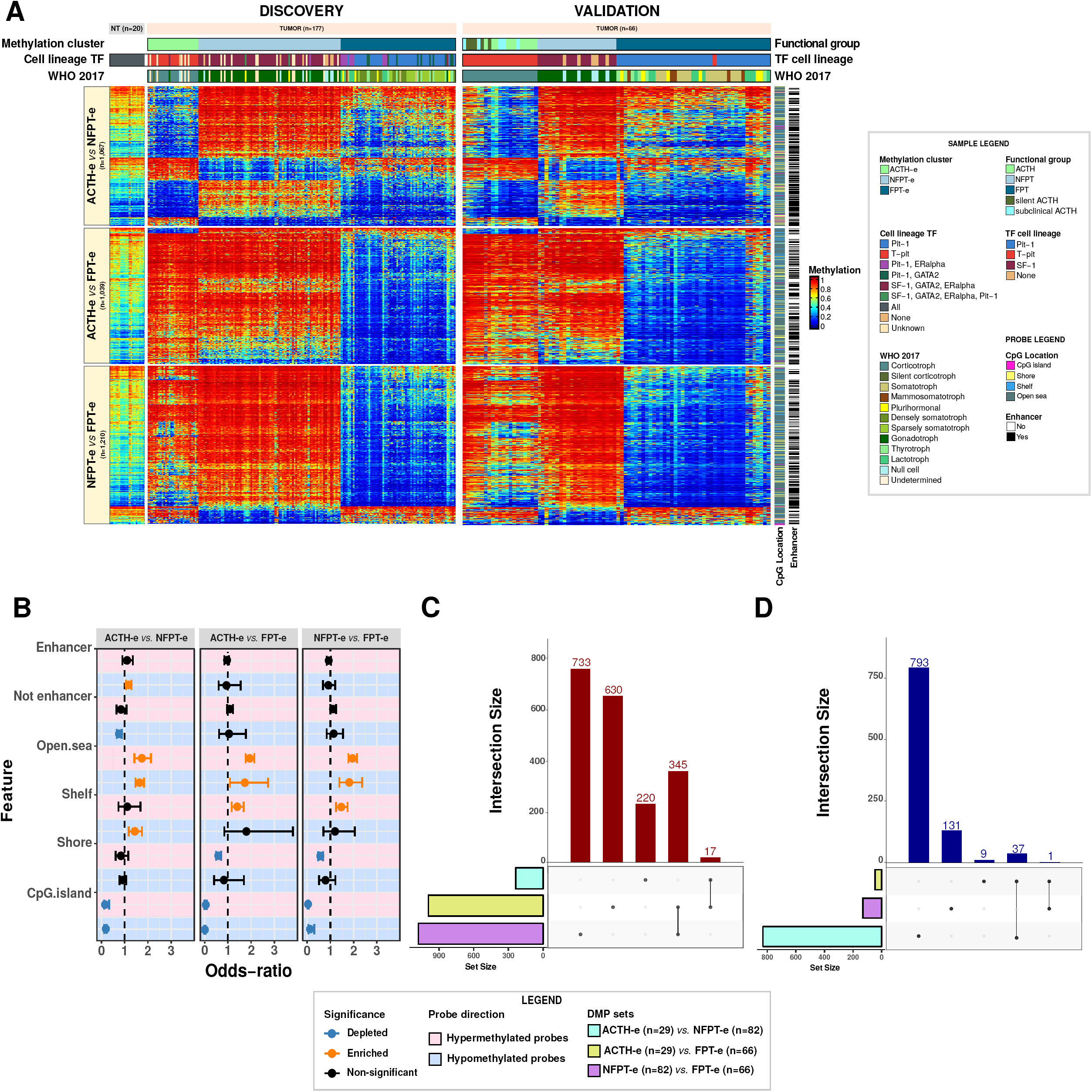
Differentially methylated distal probes define three clusters. **A)** Heatmap of the differentially methylated CpG probes (DMPs) from pairwise comparisons among clusters. Heatmap on the left displays the methylation landscape of the pan-pit cohort (discovery, n=177) and the heatmap on the right depicts the methylation patterns of the corresponding CpG applied to the Neou cohort (validation, n=86). **(B)** Odds Ratio for the frequencies of DMPs for each pairwise comparison, depicting genomic location and enhancer overlap relative to the expected genome-wide distribution of 450K probes. Relative to the first cluster of the pairwise comparison label, red- and blue-shaded boxes represent hypermethylated and hypomethylated probes respectively. **(C)** Barplots showing the number of hypermethylated DMP that are unique or common in the intersection among the three differentially methylated probe sets. **(D)** Barplots showing the number of hypomethylated DMP that are unique or common in the intersection among the three differentially methylated probe sets.

When we surveyed for DNA motifs in differentially methylated enhancer candidates that overlapped our DMPs, we identified 31 TF predicted to bind to DNA motifs in the 295 hypomethylated eDMP from the comparison between ACTH-e *vs.* NFPT-e (adjusted p-value<0.05). The top binding site was predicted to bind to Fra1/Fra2 subunits of the Activator protein (AP-1) (Figure 3A). Hypermethylated eDMP (n=88) predicted to bind to NR5A2 presented a trend to be significant (adjusted p-value= 0.06). Hypermethylated DMP from ACTH-e *vs.* FPT-e comprised cluster-related hypermethylated eDMP (n=360) presented Basic -Leucine Zipper domain (bZIP) motifs predicted to bind to 17 TFs, which are also known to interact with AP-1 subunits (Figure 3A). NFPT-e *vs.* FPT-e comparison retrieved hypermethylated eDMP-related DNA motifs (n=398) predicted to bind to 12 TF. The top most enriched TF was Jun, another subunit of AP-1 and related to the bZIP DNA binding proteins (Figure 3A). No adjusted significant DNA motifs were retrieved from the hypomethylated eDMP regions from ACTH-e *vs.* FPT-e or from NFPT-e *vs.* FPT-e comparisons (n=15 or 48 hypo-DMP, respectively). The eDMPs related to the top three high-scored enhancers are depicted in Figure 3B. Their respective associated target gene (TEAD3, PTCH1 and BAHCC1) expression levels are depicted in Figure 3C.

**Figure 3.**
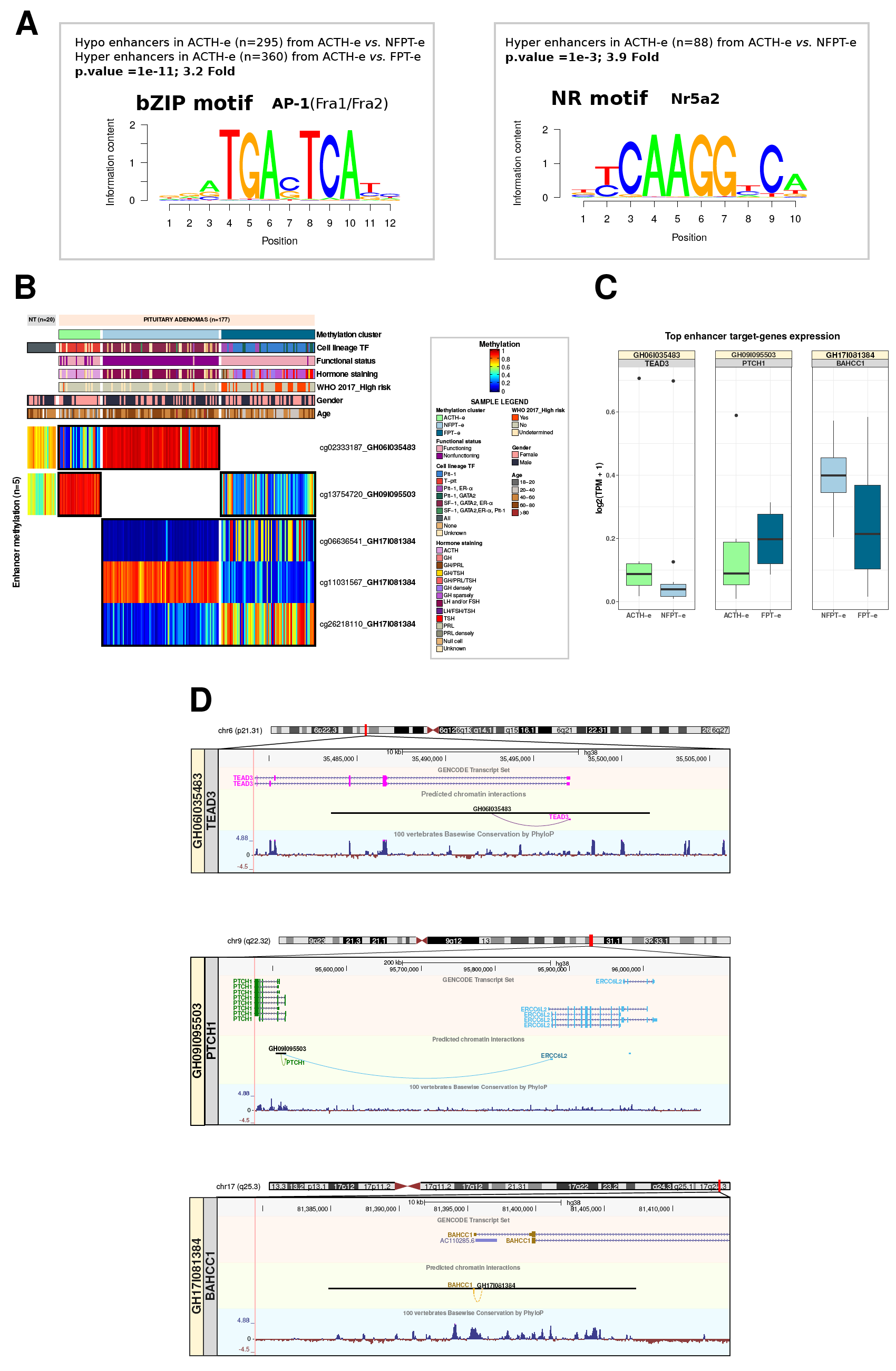
Differentially methylated candidate enhancers predicted AP-1 and NR5A2 transcription factors and respective putative enhancer-target genes. **(A)** Representation of DNA motifs significantly enriched among differentially methylated enhancer regions obtained from the pairwise comparison between ACTH-e, NFPT-e and FPT-e. **(B)** Methylation of probes in the top three highest-scored enhancers. **(C)** Expression of the high scored enhancer-target genes expression, per methylation cluster. **(D)** Genome browser view of top enhancers and the highest scored target genes, depicting the GENCODE transcripts, the predicted chromatin interactions and the conservation among vertebrates.

We observed a tendency of a negative association between the methylation degree of the eDMP and the expression of some of their putative target genes (p>0.05). For instance, in association with the correspondent hypomethylation in the eDMP, the expression of TEAD3 was higher in ACTH-e than in the NFPT-e cluster (Figure 3C). Hypomethylated eDMP in the FPT-e cluster (compared to ACTH-e) was related to upregulation of PTCH1 (Figure 3B-C) and ERCC6L2 target genes (not shown) in the FPT-e cluster. In the comparison between NFPT-e and FPT-e, we identified a cluster of 3 CpG probes overlapping a large enhancer (GH171061384) that presented a heterogeneous methylation pattern. Two out of three probes in this eDMP were hypomethylated in NFPT-e in relation to FPT-e. Their putative highest scored target genes, BAHCC1 and a lncRNA (AC110285.7), were upregulated in NFPT-e. A Genome Browser overview is used to highlight chromatin interactions between the top enhancer and putative genes pairs (Figure 3D).

### 3.3 Pituitary-related transcription factors Tpit (TBX19) and SF1 (NR5A1)-related probes differentiate specific methylation clusters

We investigated whether pituitary-specific TF probes were differentially methylated among the clusters. We found 125 probes-related to these TF (data not shown). Among the pituitary-specific TF investigated in our analysis, SF1 (NR5A1) and Tpit (TBX19) were significantly differentially methylated between two or three clusters (Figure 4A). SF1-related DMPs were all located within enhancer intronic regions (n=2), while Tpit-related DMPs mainly overlapped promoter (n=3) and intronic regions (n=2). SF1 DMPs were significantly hypermethylated in ACTH-e and FPT-e when compared to the NFPT-e (Figure 4A). Tpit-DMPs were entirely hypomethylated in ACTH-e in comparison with NFPT-e (promoter/intronic) and significantly distinguished from FPT-e-DMPs by hypomethylation in the promoter region (Figure 4B). The intronic Tpit region also distinguished the hypermethylated NFPT-e from the hypomethylated FPT-e cluster (Figure 4A). Taken together, T-pit methylation significantly distinguished all three methylation clusters; SF1 intronic methylation distinguished ACTH-e and NFPT-e and, less significantly, NFPT-e from FPT-e cluster. We established SF-1-related cutoffs for the two probes that significantly distinguished ACTH-e and NFPT-e. Specimens that presented SF1-related probes with methylation values below 0.5 (cg14143574) and 0.3 (cg02853418) were assigned to belong to NFPT-e, while beta-values above those cutoffs were considered to belong to ACTH-e cluster (Figure 4B). The application of these cutoffs to the Neou cohort correctly assigned 95% to either ACTH/T-pit or NFPT/ SF1 specimens (χ^2^-squared p-value = 1.68e-08). Likewise, three out of five T-pit-related DMP accurately distinguished ACTH-e from non-ACTH-e specimens (NFPT-e and FPT-e). Specimens with CpG methylation below the thresholds for at least two out of those 3 probes, *i.e.* 0.5 (cg26160839), 0.7 (cg01732037) and 0.5 (cg15140722), assigned samples to the ACTH-e group (Figure 4B). Applying this criteria to the validation cohort ^10^, 75% of the ACTH tumors were classified accordingly while 100% of the other functioning and nonfunctioning PT were assigned as the nonACTH category (χ^2^-squared p-value = 7.56 e-14).

**Figure 4.**
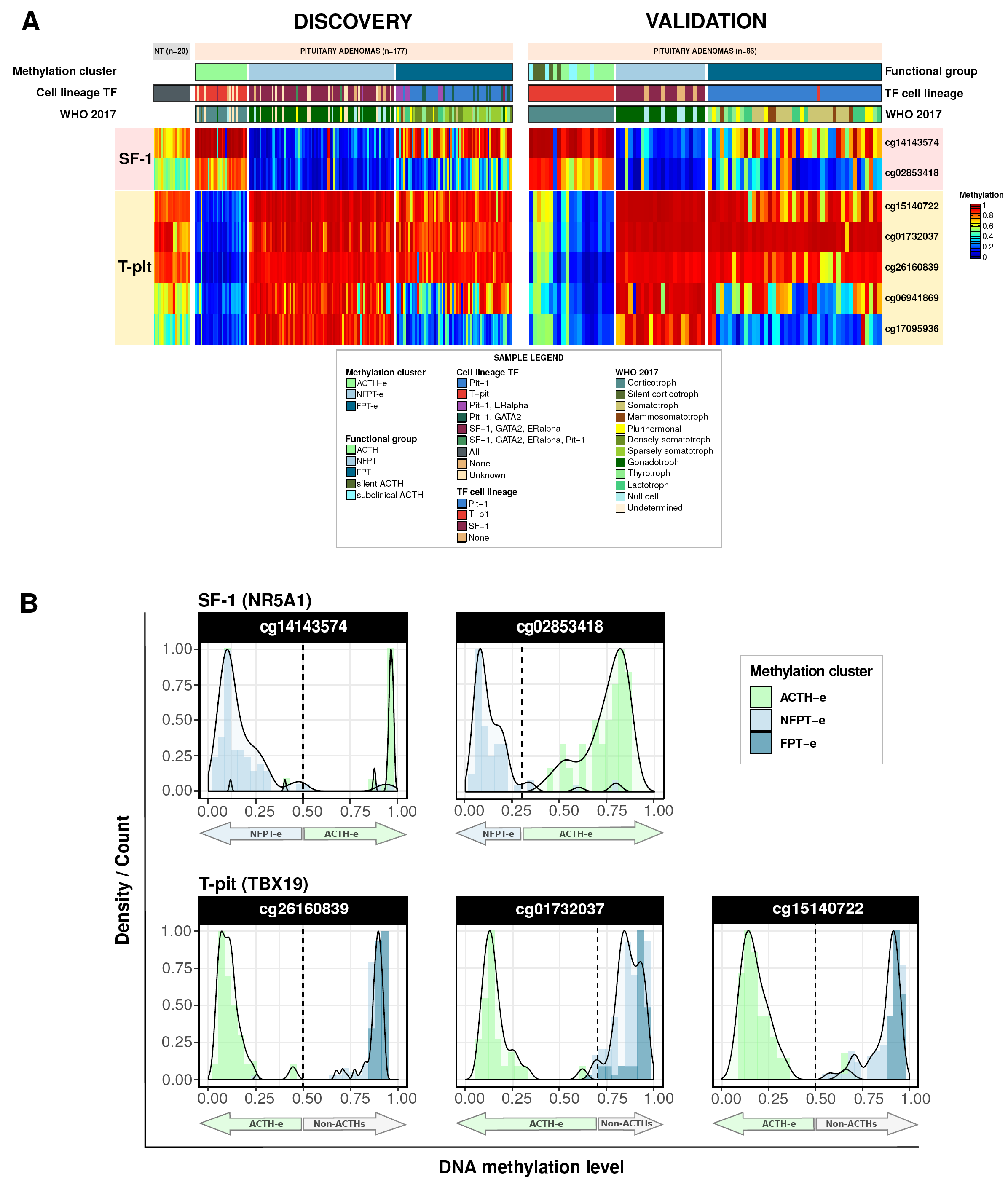
Pituitary-specific transcription factors found differentially methylated among the three methylation clusters. **(A)** Heatmap indicating significant differentially methylated probes associated with SF-1 (NR5A1) and T-pit (TBX19) among the three methylation clusters: ACTH-e, NFPT-e and FPT-e. Heatmap on the left displays the pan-pit cohort (discovery, n=177) and the heatmap on the right depicts the methylation patterns of the corresponding CpG applied to the Neou cohort (validation, n=86). **(B)** Density plots on the distribution of beta value and tentative thresholds (vertical dashed lines) for SF-1- and T-pit-related differentially methylated probes that distinguish the ACTH-e from NFPTs and ACTH from non-ACTHs, respectively.

## 4 Discussion

We conducted a comprehensive and integrative analysis between the molecular and clinicopathological features of the largest multicentric methylome cohort of pituitary gland specimens to date. An unsupervised analysis of the methylome cohort defined three groups of PT characterized by hypo-, hyper- or intermediately methylated profiles mainly segregated by functionality (functioning and nonfunctioning) as well as adenohypophyseal cell lineages, namely FPT-e, NFPT-e and ACTH-e, respectively, similar to the results reported in recent studies ^10,15^. These methylation clusters contain subtypes that recall and provide an epigenetic support for the pituitary cell lineages-based criteria to classify PT recommended by the 2017 WHO classification, particularly to the Pit-1-derived tumors ^5^.

Notably, in the NFPT-e cluster, the methylome similarities between gonadotrophs and null cell adenomas reported by us and others ^10,15,28^ may indicate that hypermethylation underlies the pathogenesis of the lack of production and/or secretion of adenohypophyseal hormones. In addition, we and others showed that silent and functioning corticotroph adenomas (i.e. overt Cushing’s disease and subclinical) shared a more similar methylation profile compared to each other and with NFPT. These results are concordant with a report that silent ACTH and gonadotroph tumors share cytopathological features and clinical behavior ^29^. Taking into account that cell-specific methylation patterns are preserved during development and tumorigenesis, it is also possible that null cell, a subset of gonadotrophs and of corticotroph adenomas share a common precursor ^30,31 10,32,33^ (Figure 1B). In addition, a study showed that a subset of functioning corticotroph adenoma displayed a more intermediate methylation profile between FPT and NFPT ^10,32,33^. Specific gene mutations such as GATA3 and USP8 accounted for the differences among these corticotroph adenomas variants ^10,28^. Altogether, these results show that corticotroph adenomas consist of a molecularly heterogeneous tumor ^10,15,28^. Whether the association between methylation patterns and mutations in these variants are drivers or passengers remains to be elucidated.

Pairwise comparisons between the methylation clusters yielded the identification of a significant number of DMP (Figure 2A) that were mainly located in enhancer regions and enriched for open sea CpGs (Figure 3B). Validating our findings, the overlap of either the DMP or the top eDMP with the methylome, assigned the specimens of an independent cohort ^10^ into similar methylation and clinicopathological groups as we observed in the pan-pit cohort. These novel results indicate that these regulatory regions may play key roles in pituitary tumorigenesis as well as in cell-specific development and differentiation as described in other tumors ^34^. The biological significance of the differentially methylated enhancers across the methylation clusters are endorsed by their association with target genes involved in pathways related to developmental cytodifferentiation and in the tumorigenic process (*e.g.* TEAD3, TAF11, lnRNA/AC110285.7, ERCC6L2, PTCH1) (Figures 3B,C)^35–42^.

Recently, the integration of multiple platforms including genomic and epigenomic features has been proposed as an unbiased molecular approach to classify PT ^10^. Despite its thoroughness, the complexity and the current cost of profiling multiple platforms to classify a sample may limit its application in the clinics. Interestingly, their methylation profiles showed similar clusters of PT as we described here. In addition the methylation-based groups were similar to the ones obtained using the pangenomic approach ^10^. Altogether, these findings provide evidence that methylome profiling *per se* may be a reliable and an unbiased approach to classify PT ^10^. In our analysis, we were able to identify specific methylation thresholds related to SF-1 and T-pit genes that significantly distinguished ACTH-e/Tpit from NFPT-e/SF1 methylation groups in our cohort that correctly assigned ACTH and NFPT specimens in the independent cohort ^10^. These results, if validated by other methods (e.g., pyrosequencing), suggest that the methylation degree of these probes could be used as an objective approach to classify PT specimens according to adenohypophyseal cell lineage.

The prognostic and predictive significance of methylation-based stratification in PT is not clear, in contrast to the findings reported in other CNS tumors such as gliomas and meningiomas ^8,43^. We and others showed that PT methylation clusters were not associated with features used as surrogates of aggressive behavior in PT (*e.g.* invasion, resistance to treatment etc) ^5,10,28^. However, the association of PT methylation clusters with a prognostic grading which has been validated as predictors of tumor aggressive behavior remains to be investigated ^8,43,44,45^.

Besides refining tumor classification and stratification, methylome profiling can also be applied to differentiate tumors originated from different tissues ^7,46^. Indeed, one study showed that distinct methylome patterns segregated PT apart from other CNS tumors^7^. Potentially, PT-specific signatures, as the ones herein described, could be helpful to distinguish PT from other primary or secondary sellar tumors whose diagnosis by morphologic and immunohistochemical approaches may be challenging and inconclusive ^7,47,48^.

In summary, we identified and validated methylation signatures that distinguished PT by distinct functional status/subtype and adenohypophyseal cell lineages. These signatures, annotated in enhancer regions, indicate the importance of these elements in the pathogenesis of these tumors. They also provide an unbiased approach to classify pituitary tumors according to the most recent classification recommended by the 2017 WHO using methylation profiling.

## Acknowledgements

The authors are grateful to the HFHS patients who consented to the usage of PT for research purposes. We thank Nancy Takacs and Heather Mengel for their administrative support; Kevin Nelson for the collection, handling and maintenance of the tumor bank at the Hermelin Brain Tumor Center; Andrea Transou for tumor pathology processing; Laura A. Hasselbach for DNA extraction; Daniel Weisenberger and team at USC Epigenome Center for assistance with DNA methylation profiling (HFHS support);Susan MacPhee for proofreading the manuscript.

## Conflict of interest

The authors declare to have no competing interests.

## Table caption

**Table 1 -** Clinicopathological classification features of pituitary tumors from a multicenter cohort. NFPT: Nonfunctioning pituitary tumor;

## Supplementary Figures

**Figure S1 -** Principal component analysis (PCA) colored by (A) cohort source; (B) by PT functional status; (C) by methylation cluster; (D) Cumulative distribution function of consensus clustering classification and (E) Consensus cluster delta area plot of consensus clustering classification.

**Figure S2 - DNA methylation-based stemness index (mDNAsi) of pan-pituitary cohort specimens, according to methylation clusters and clinicopathological features.**

**(A)** By methylation clusters.

**(B)** By functional status.

**(C)** By the high risk subtypes according to the 2017 WHO classification.

**(D)** By gender.

**(E)** By age.

*=p<0.05;**=p<0.01;***=p<0.001;****=p<0.0001

**Figure S3 - Supervised analysis of the methylome according to invasion status.**

**(A)** Heatmap of the 13 differentially methylated probes between invasive *vs.* noninvasive PT (n=34 *vs.* n=46, respectively; p< 0.001 and Δβ=0.15).

**(B)** Venn diagram depicting common and distinct differentially methylated genes from our study compared to several studies.

**(C)** Differentially methylated probes overlapping enhancer regions (eDMPs) and the respective candidate enhancer-target genes correlation according to genomic context.

**(D)** Differentially methylated probes overlapping enhancer regions (eDMPs) and the respective candidate enhancer-target genes correlation according to invasion status.

**Figure S4 - DNA methylation-based stemness index and RNA-based stemness index of Pan-pit cohort and integrative subset cohort based on the invasion status.**

**(A)** DNA methylation-based stemness index (mDNAsi) of the pan-pituitary specimens cohort colored by invasion status (n=197).

**(B)** DNA methylation-based stemness index (mDNAsi) of a subset of the pan-pituitary cohort colored by invasion status (n=23)

**(C)** RNA expression-based stemness index (mRNAsi) of a PT cohort subset, colored by invasion status (n=23).

## Supplementary tables

**Table S1 -** Compiled clinical information for each patient pituitary specimen.

**Table S2 -** Differentially methylated probes from supervised comparisons. 1-3) differentially methylated probes (DMP) among the three methylation clusters comparisons; 4) DMPs between tumorous and non-tumorous pituitary specimens. 5) DMPs between invasive and noninvasive PT.

## Notes

**Funding:** This work was supported by the Henry Ford Health System, Department of Neurosurgery, and the Hermelin Brain Tumor Center. Additionally, MSM and MC are supported by The São Paulo Research Foundation (FAPES), Brazil (#16/11039-3; #17/10357-4,#14/03989-6); AC and KPA by Henry Ford Hospital (A30935, A30957; GME 202199); LMP, HN, AD, MW, and AM are supported by the National Institutes of Health (R01CA222146), HN, TSS, TMM, LMP, and AD are supported by the Department of Defense (CA170278).

### Competing Interest Statement

The authors have declared no competing interest.

