## Supplementary figures and images for "DNA Methylation-based Signatures Classify Sporadic Pituitary Tumors According to Clinicopathological Features"

### Supplementary Figure S1

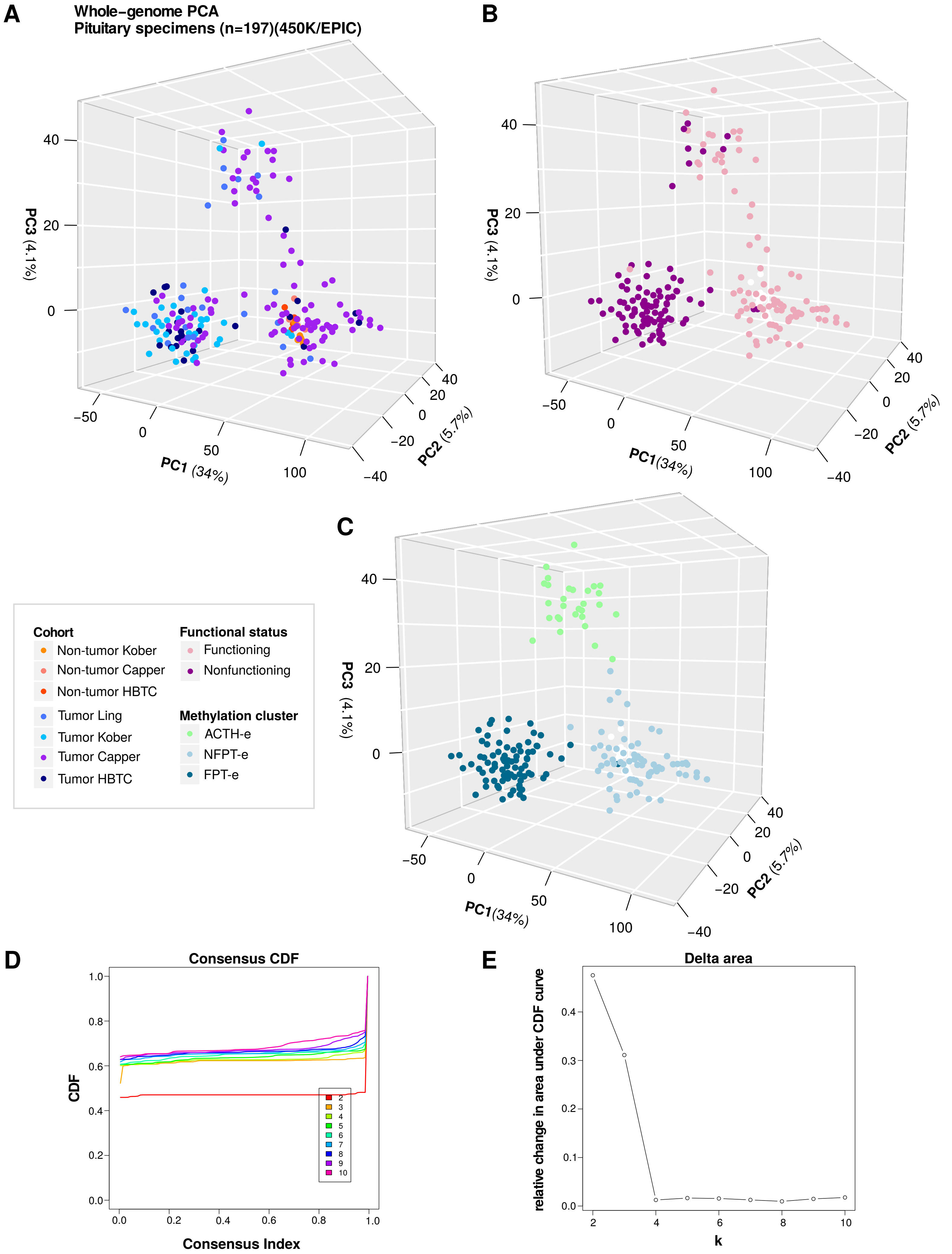

### Supplementary Figure S2

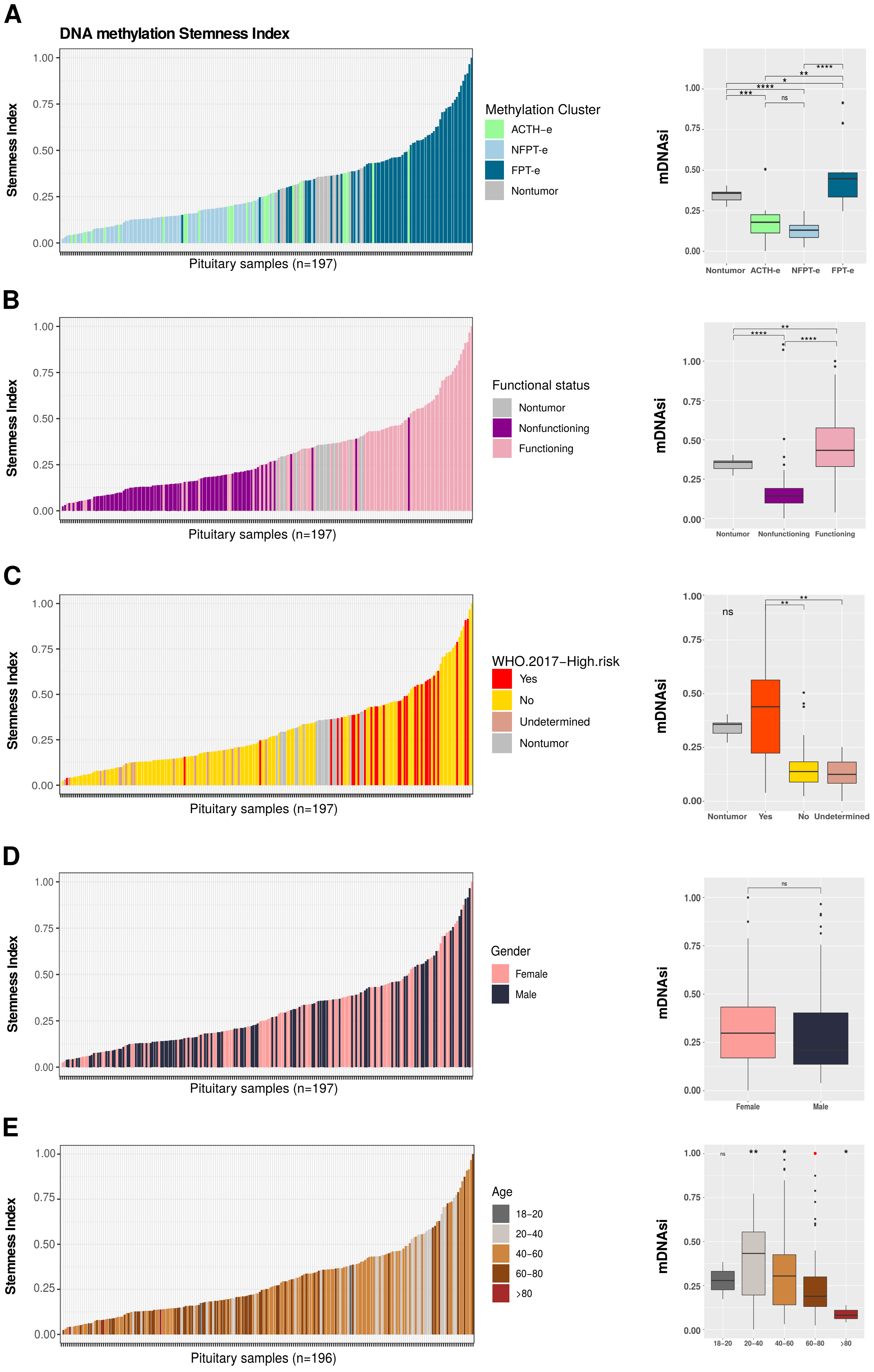

### Supplementary Figure S3

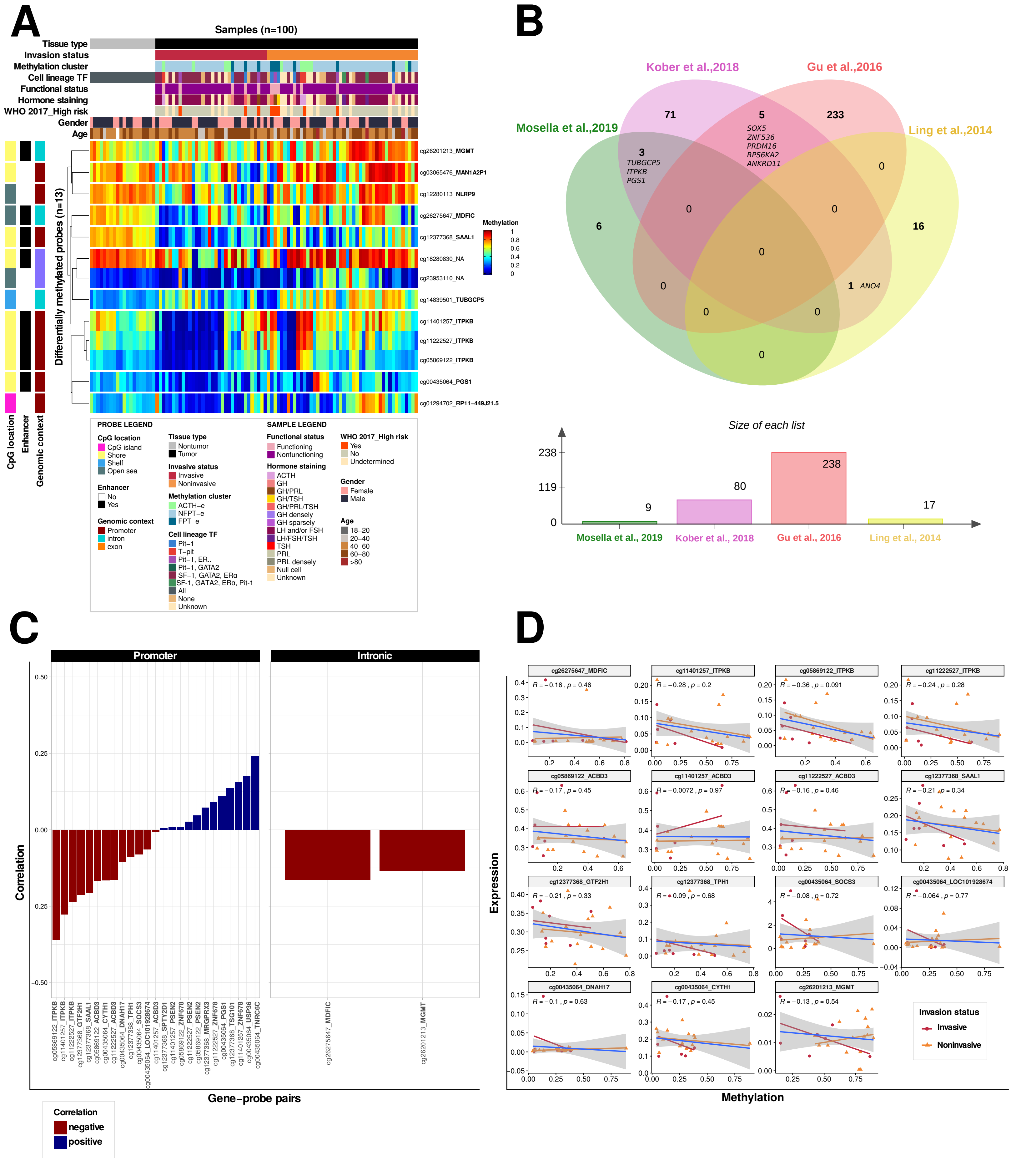

### Supplementary Figure S4

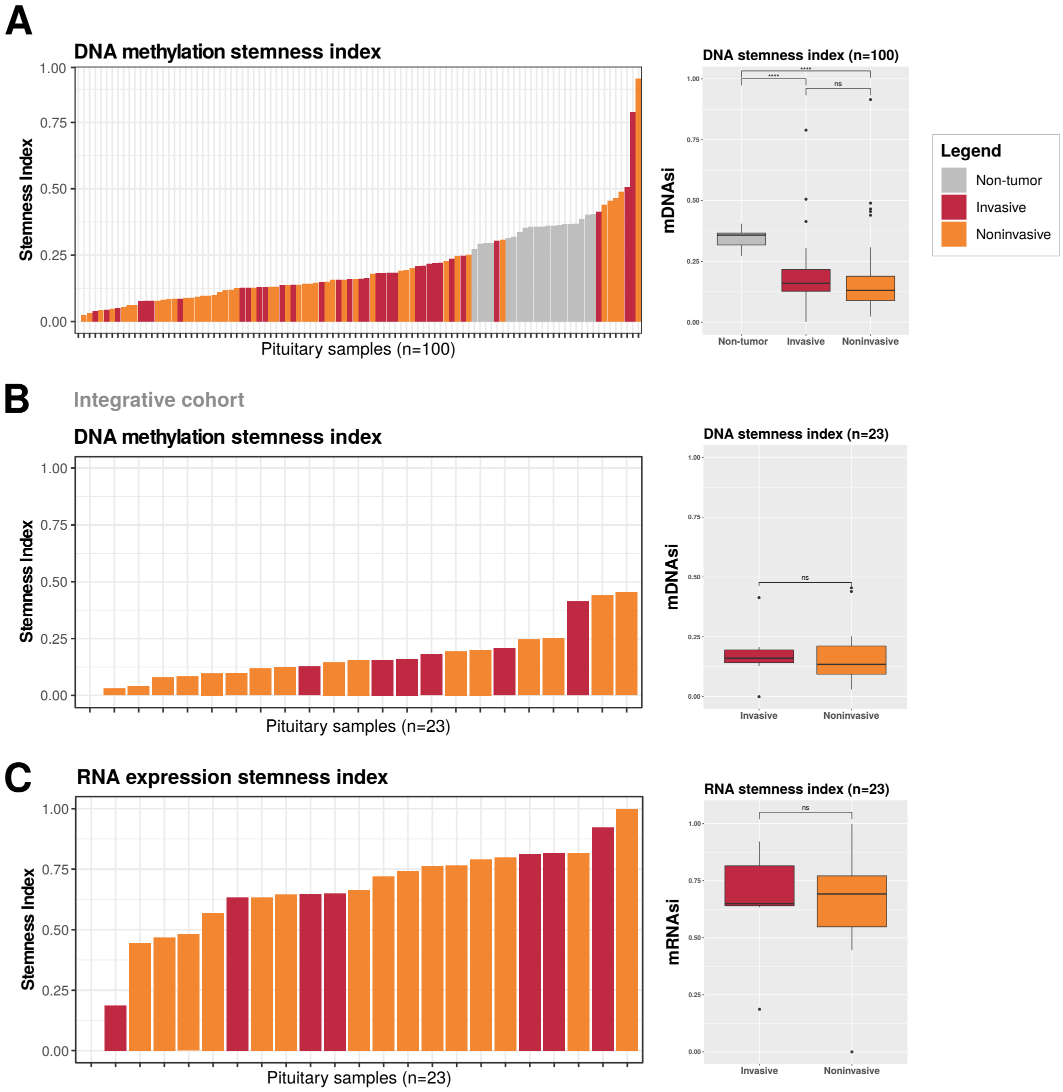
